# Bundling and segregation affects “liking”, but not “wanting”, in an insect

**DOI:** 10.1101/2022.05.24.493357

**Authors:** Massimo De Agrò, Chiara Matschunas, Tomer J. Czaczkes

## Abstract

Behavioural economists have identified many psychological manipulations which affect perceived value, although value in humans is not a unitary experience, with “liking” and “wanting” being neurologically separate processes. A prominent example of this is bundling, in which several small gains (or costs) are experienced as more valuable (or costly) than if the same total amount is presented together. While extensively demonstrated in humans, to our knowledge this effect has never been investigated in an animal, let alone an invertebrate. We trained individual *Lasius niger* workers to two of three conditions in which either costs (travel distance), gains (sucrose reward), or both were either bundled or segregated: A) both costs and gains bundled, B) both segregated, and C) only gains segregated. We recorded pheromone deposition on the ants’ return trips to the nest as measure of “liking”. After training, we offer the ants a binary choice between odours associated with the treatments, as a measure of “wanting”. While bundling treatment did not affect choice, i.e. “wanting”, it strongly influenced pheromone deposition, i.e. “liking”. Ants deposited c. 80% more pheromone when rewards were segregated but costs bundled as compared with both costs and rewards being bundled. This pattern is further complicated by the pairwise experience each animal made, and which of the treatments it experiences first during training. The current study is the first to demonstrate a bundling effect in an animal, and the first to report a dichotomy between “liking” and “wanting” in an insect. We propose that the deviation between “wanting” and “liking” in this case is due to the unique nature of distance perception in ants, which is recorded linearly, while almost all other sensory perception in animals is logarithmic.

## Introduction

Broadly speaking, when an animal must choose between options, it can employ one of three different strategies, characterized by different levels of precision: random choice; following a heuristic or rule of thumb; or comparison of outcome value and choice for the highest. Traditional economic theory, exemplified by Expected Utility Theory, assumes (human) decision-makers are rational and perform strict value-based choice (Mankiw, 2011). The now well-established field of behavioural economics has vigorously pushed back against this idea, demonstrating that humans often make decisions which are not fully logical, economically rational, or “optimal” (Camerer et al., 2011; Tversky and Kahneman, 1974), even when actively comparing options.

A major insight lying at the heart of behavioural economics is that value is *perceived*. This can lead directly to deviations from optimality: between the acquisition of the information and the evaluation of possible outcomes, something gets lost in translation. As has been well established for over a century by the study of pyschophysics, perception is non-linear, usually on a logarithmic scale (Gescheider, 1997). Value perception for humans is likewise nonlinear, as famously stated by Kahneman and Tversky, and extensively demonstrated thereafter (Camerer, 2004; Kahneman and Tversky, 1979). Moreover, humans weigh losses more strongly than gains. Finally, value is relative, usually to an expectation or some sort of anchor, see Figure 1.

**Figure 1:**
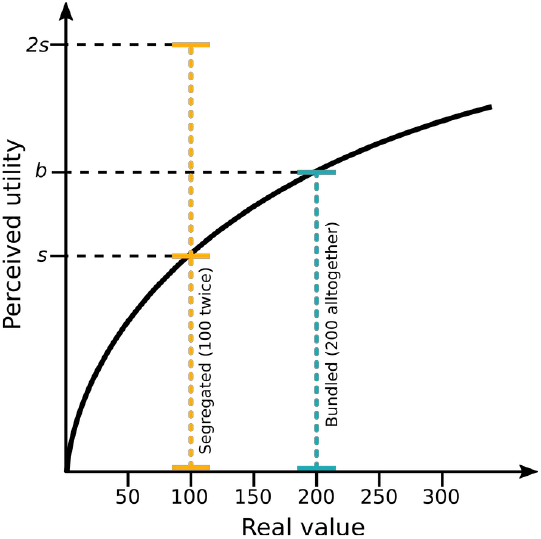
Simplified schematic of Prospect Theory from Kahneman & Tversky (1979), with a graphical illustration of bundling and segregation. On the x-axis, actual value of a gain or a loss (here exemplified with money). Perceived utility does not scales linearly with value, but logartitmically. Receiving a gain of €100 (segregated) twice will produce a level of “happiness” of “2s”, more than the level “b” perceived when receiving €200 all together (Bundled). The same is true for “losing” the same amounts, where two losses of €100 are felt stronger than a single one of €200.

The non-linear nature of perceived value results in many behavioural biases, a very prominent one being the bundling verses segregation effect. Crucial to the current experiment, the perceived value of a compound item or option can be changed by presenting it either as one option (bundling), or as multiple small parts (segregation). Due to diminishing returns, bundling results in a weaker total sensation than segregation– either lower value for positive value options, or lower cost for negative value options (see Figure 1). This is because more of the sensation occurs on the shallow part of the curve. This fact is regularly exploited by consumer psychologists and marketing experts, for example by bundling option when selling new cars: when spending €50,000 on a new car, spending €51,000 for the model with included sound system might not be experienced as painfully costly, even if, considered by itself, €1000 might be more than most people are willing to spend on a sound system for a car. Bundling and segregation have been extensively studied by consumer psychologists (Johnson et al., 1999; Naylor and Frank, 2001; Noone and Mattila, 2009).

As in the study of human economics, non-human animals have been treated and modelled as rational economic agents, leading to deep insights into animal behaviour via the Optimal Foraging Theory framework (Davies et al., 2012; Emlen, 1966; MacArthur and Pianka, 1966; Pyke et al., 1977). However, often inspired directly by behavioural economic research on humans, predictable deviations from optimality and rationality have been described (Zentall, 2015). For example, pigeons, rats, and ants all show a preference for high-effort over low-effort associated options (Clement et al., 2000; Czaczkes et al., 2018a; Lydall et al., 2010), much as humans do (Norton et al., 2012). Similarly, much as in humans, the addition of an irrelevant option in an option set (a “decoy”) can change the preference structure in many animals, including birds, cats, bees, and ants (Bateson et al., 2002, 2003; Parrish et al., 2015a; Sasaki and Pratt, 2011; Scarpi, 2011; Schuck-Paim et al., 2004; Shafir et al., 2002).

Ants and bees also show relative value perception, changing the perceived value of an option depending on their expectations (Bitterman, 1976; Couvillon and Bitterman, 1984; Wendt et al., 2019; Wendt and Czaczkes, 2020). An ant which is trained to expect a very sweet reward, for example, will be more likely to reject a moderate reward than an ant which was expecting a moderate one – a negative contrast. Likewise, ants expecting a very mild reward are more likely to accept a moderate reward than ants which were expecting that quality - a positive contrast (Wendt et al., 2019). These changes are also mirrored in the ants’ deposition of recruitment pheromone: ants deposit more pheromone to resources they perceive as higher quality (Beckers et al., 1993; Jackson and Châline, 2007), and indeed ants deposit more pheromone for moderate rewards if they had been expected poor quality, and less pheromone if they were expecting high quality. This demonstrates that a key aspect of Prospect Theory - relative value perception – is present in insects as well as humans.

Crucially for the current experiment, there is evidence that social insects, especially ants, perceive value logarithmically. Wendt et. al. (2019) demonstrated a faster rise in food acceptance in the lower range of acceptance food qualities than in the higher range. Recently, De Agrò et al. (2021) demonstrated that *Lasius niger* ants have a strong aversion to risky food sources (i.e. with fluctuating quality), which could be fully explained by logarithmic value perception, as predicted by Prospect Theory. Ants prefer a certain food source offering 0.55M sucrose to one which fluctuates between 0.1 and 1.0, but if the options are logarithmically balanced (0.3M vs 0.1 or 0.9), ants are completely indifferent.

While the bundling versus segregation effect is extremely well studied in humans, surprisingly, to our knowledge no attempt has been made to examine it in animals. In a related study, chimps were shown to prefer whole rewards (potato chips) to rewards which were broken into smaller pieces, even when the broken rewards had an higher absolute quantity (Parrish et al., 2015b). However, consumption of the whole and broken rewards took the same amount of time, and rewards were chosen before consumption could begin, so it is likely that both broken and whole rewards were considered as one unitary bundle.

The current study is the first to examine bundling verses segregation in animal, using the black garden ant, *Lasius niger*, as a model. We take a dual approach, quantifying both perceived value by recording the amount of recruitment pheromone deposited directly after encountering a reward, and relative preference, by offering trained ants a choice between bundled and segregated options. We routinely collect pheromone deposition data in order to make inferences about how individual level behaviour may interact with group level behaviour. However, the results of the current study (see results and discussion) caused us realise that pheromone deposition behaviour may reflect something fundamentally different to binary choice assays. While binary choices assays inform us about which option the animal “wants”, pheromone deposition informs us about how much the animal has “liked” the reward. The duality between “liking” and “wanting”, first described in rats (Berridge et al., 1989) and then confirmed extensively in humans (Brauer and De Wit, 1997; Leyton et al., 2002; Pool et al., 2015), distinguishes two aspects of reward. “Liking” refers to acutely perceived hedonic reactions, while “wanting” refers to motivation and desire for a goal – the quotation marks distinguish these from the conscious, cognitive forms of enjoyment and desire that the words imply in everyday conversation. While “liking” and “wanting” usually co-vary, these are neurologically separate processes, and can, in vertebrates, be separately inactivated (Berridge et al., 1989) or enhanced (Berridge and Valenstein, 1991; Leyton et al., 2002; Treit and Berridge, 1990). Hyperactivity of “wanting” but not “liking” is strongly linked to addiction and eating disorders (Berridge and Robinson, 2016; Finlayson et al., 2007; Robinson et al., 2016).While the need for a method of distinguishing “liking” and “wanting” in insects has been highlighted (Søvik et al., 2015), to our knowledge the ant pheromone deposition paradigm we propose is the first concrete approach to measuring these separately.

In order to study bundling and segregation in ants, we segregated rewards spatially, using small sucrose drops along a linear runway. However, while this segregates rewards, it also segregates a potential cost – the walking distance to the reward. Thus, we first demonstrate that ants do indeed prefer closer to farther food sources, and thus that food distance is considered a cost. We then train individual ants to two of three treatments: rewards and costs bundled (‘Bundled’), rewards and costs segregated (‘segregated all’), and rewards segregated but costs bundled (‘segregated reward’). We record pheromone depositions on the ant’s return from each treatment type (= “liking”), and then ask ants to choose between the pair of treatments they were trained on (= “wanting”). We predicted that the ‘segregated reward’s treatment would be the most “liked” and “wanted” option, as bundling of costs should minimise their impact, and segregation of rewards should boost theirs. As we did not know the relative strength of the rewards and costs, we had no strong a priori predictions about the relative perception of ‘Bundled’ and ‘Segregated All’. However, as Prospect Theory predicts that losses are weighted more strongly than gains (Kahneman and Tversky, 1979), we had a weak expectation that ‘Bundled’ would be perceived as slightly better than ‘all segregated’.

## Materials and methods

### Subjects

We used 19 queenless *Lasius niger* colony fragments, consisting of around 1000 ants each. Each fragment was collected from a different wild colony on the University of Regensburg campus. Workers from colony fragments forage, deposit pheromone and learn well (Evison et al., 2008; Oberhauser et al., 2018). Each fragment was housed in a transparent plastic box (30×20×40cm), with a layer of plaster on the bottom. A circular plaster nest, 14cm in diameter and 2 cm thick, was also provided. The colonies were kept at room temperature (21-25 °C) and humidity (45-55%), on 12:12 light:dark cycle. Each colony was fed exclusively on 0.5M sucrose solution ad libitum, and deprived of food 4 days prior to each test. Water was provided ad libitum and was always present.

### Procedure

All four experiments reported in this paper used a conditioning procedure as described in Czaczkes (2018). The procedure was generally the same, with a few modifications dictated by the specific conditions.

For each tested subject the procedure started by connecting a drawbridge to the nest box. This bridge was composed of a 20cm long, 1cm wide, slanted section, one end of which laid on the plaster floor of the nest box. The other end led to a straight 10cm long, 1cm wide, runway section. Both of these were covered by unscented paper overlays. Depending on the visit, this bridge could lead either to a straight runway, or to the stem of a Y-maze. In the first visit, multiple ants were allowed on the bridge. A 0.5M sucrose solution drop was placed at the end of the bridge. The first ant to reach the drop and start drinking was marked on the abdomen with a dot of acrylic paint. The non-marked ants were gently returned to the nest box, while the marked one was allowed to drink to satiation, then allowed to return to the nest on her own. In the nest the marked ant performed trophallaxis with nest-mates, and then returned to the bridge location ready for the following visit, which varied depending on the experiment being run.

### Pilot Experiment – Is increased food distance negatively perceived?

As described in the introduction, segregated rewards should be perceived as being of higher value than an equal-quality bundled alternative. However, this is also true for punishments. Since in our experiment we segregated rewards by placing them on different parts of a long runway (see next paragraph), we needed to test whether increased travelled distance makes rewards less preferred.

The previously-marked ant was allowed onto the bridge. This time, the bridge was attached to either a 25cm or 75cm long runway (systematically varied). This runway was covered with a scented paper overlay (either rose or lemon odour). Scenting was achieved by storing the overlays in a sealed plastic box with 3 drops of food flavouring for at least 24 hours. At the end of the runway, we placed a high-quality (1.5M) sucrose solution drop, flavoured with the same smell as the runway (at a ratio of 1μl food flavour per ml sucrose solution). The ant eventually found the drop, drank to satiation, and then went back to the nest. At this point, we discarded the scented overlays, in order to remove the deposited pheromone.

As soon as the ant unloaded, it was allowed back onto the bridge again. The bridge now connected to the reciprocal runway length (25cm if the previous visit was to 75, and vice versa). This runway was covered with paper scented with a different smell to the previous visit. At the end of the runway, the ant again found a drop of 1.5M sucrose solution, again flavoured to match the paper overlay. After drinking, the ant again allowed to return to the nest.

The same procedure was repeated another time. Thus, the ant experienced the long and the short runways twice each. For all visits, we measured the number of times the ant deposited pheromone on the scented runway, both the way towards the drop and the way back. Pheromone deposition in *Lasius niger* is a stereotyped behaviour, and easily quantified by eye (Beckers et al., 1993).

After these 4 visits, the ant was allowed onto the bridge one last time. This time, the bridge was connected to the 10cm long stem of a Y-maze. The stem was covered with unscented paper, and tapered to a 2mm wide point. Here, the two arms of the Y-maze started, also tapered, in order to ensure that the ant to contacted both arms at the same time once at the end of the stem. One of the two arms was scented with the long runway odour, while the other was scented with the short runway odour. We noted on which of the two arms the ant ran for at least 2cm (considered the ‘initial’ decision), and at the end of which of the two it arrived first (considered the ‘final’ decision). Once this happened, the ant was picked up with a piece of paper and moved back at the start of the stem to be retested. This way, we could test the ants’ preference three times. After the test, the ant was permanently removed from the colony.

A total of 24 ants from 5 different colonies was used for this experiment.

### Main experiment – Bundling vs Segregation

Having established that increased travel distance makes food sources less attractive (see results), we proceeded with the main experiment.

The procedure is very similar to the pilot. Each marked ant was allowed onto the bridge, after which it was presented with a scented runway. This scented runway could correspond to one of three different treatments:

#### A) ‘Segregated All’ (rewards and costs) (Fig. 2A)

The ant encountered a 75cm long runway. Every 25cm, the runway tapered to 2mm in width with a 0.2μl, 1.5M, sucrose solution drop on the taper. After the drop, the runway widened again, until the next 25cm taper. Here the ant found a second drop, identical to the previously encountered one. The runway proceeded for a third 25cm section, ending in a large drop. To avoid evaporation, as well as limiting the risk of the ant bypassing the rewards without noticing, the drops were delivered with a micropipette when the ant reached the designated position, rather than being placed prior to testing.

**Figure 2:**
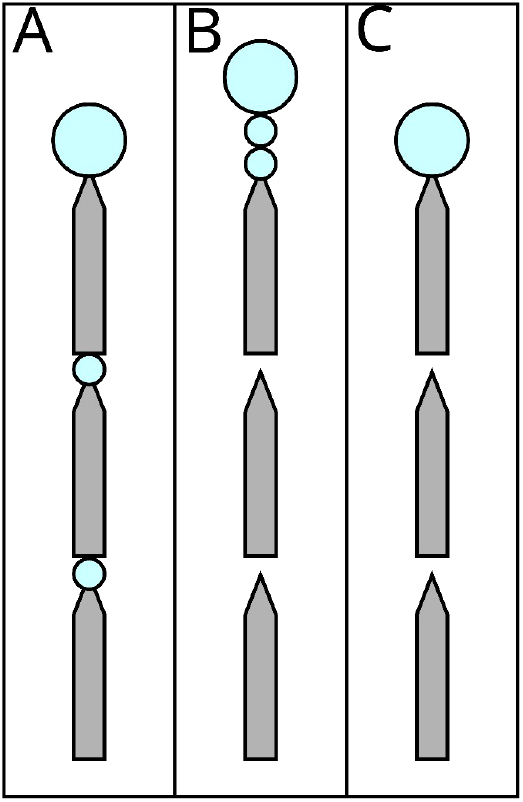
The three possible experimental treatments. Grey shapes represent the runway segments, each 25cm long, 1cm wide, tapering to 2mm to ensure that ants encounter the sucrose drops. Big blue circles represent ad-libitum 1.5M sucrose solution, small circles represent 0.2μl drops which the ants can drink, but will not satiate them. In A) ‘Segregated All”, both the costs (travel over the runways) and the rewards (drops of 1.5M sucrose, blue circles) are segregated. In B) ‘Segregated Rewards”, only the rewards are segregated. In C) ‘Bundled’ both the costs and the rewards are bundled.

In this treatment, the reward (drops) was segregated into 3 different experiences, and as such its combined value should be perceived as higher than one being presented as a single one. The volume of the first two drops was selected in order to ensure that the ant would reach the third drop without becoming satiated after the first or the second one: The crop volume of *L. niger* foragers is under 1μl (Mailleux et al., 2000), and ants which encounter such drops drink them and then continue walking forwards (Czaczkes et al., 2019). The last drop instead was much larger, to ensure that ants could drink to satiation, and thus avoiding other possible discounting effects, such as disliking not being completely satiated (Mailleux, 2006; Mailleux et al., 2005).

However, in this treatment also the cost is segregated: Rather than being experienced as a single 75cm long runway, the ant encountered three 25cm ones, interspersed by rewards. Thus, this condition is expected to enhance both the perceived value and the perceived cost.

#### B) ‘Segregated Reward’ (bundled costs) (Fig. 2B)

In this second treatment, the ant encountered a 75cm long runway. At every 25cm mark, a narrowing portion was present, but in this treatment no small sugar drops were provided. The narrowing of the paper was maintained to ensure consistency with the previous treatment. To assure consistency among treatments, the experimenter followed the same procedure of the ‘segregated all condition’: the micropipette was brought to the narrowing point at the end of the 25cm runway when the ant got near it and the plunger depressed delivering no drop (i.e. a sham treatment). At the end of the 75cm, two 0.2ul, 1.5M drops were presented in short succession, just 5mm from each other. After another 5mm, the third ad libitum drop was placed. Thus, in this treatment, the distance travelled (=cost) remained bundled, while the reward was still segregated.

#### C) ‘Bundled’ (rewards and costs) (Fig. 2C)

In this third treatment, the ants encountered the same runway as above. However, instead of presenting three drops at the end, only the third ad libitum drop was offered. Thus, in this treatment, both the reward and the cost are bundled. The sham pipetting was also carried out in this treatment.

#### Pairwise training and a priori hypotheses

We performed three different conditions, corresponding to the three different pairings of the three treatments (A vs B, A vs C, B vs C). 40 ants were tested per condition, for a total of 120. Each ant would experience one of the three treatments for the first visit, associated with a distinct odour. On the subsequent one, the animal encountered a second treatment, associated with the another odour. This procedure was repeated for 4 times, for a total of 8 visits alternating between the two selected treatments. In the end, the ant was presented with a Y-maze, and had to choose between the two treatment odours.

Broadly, we expect the following preference structure:

- In condition 1 (B vs C), the ‘Segregated Reward’ treatment should be preferred over the ‘Bundled’ one. This is expected due to the introduced bundling effect.
- In condition 2 (B vs A), the ‘Segregated Reward’ treatment should be preferred over the ‘Segregated All’ one. This is expected as the boosting of reward by segregation effect is identical, while the ‘Segregated All’ condition also boost the cost.
- In condition 3 (A vs C), the ‘Bundled’ treatment should be preferred over the ‘Segregated All’ one. This is because costs are often more heavily weighted than gains (Lakshminarayanan et al., 2011; Tversky and Kahneman, 1981), and so boosting both rewards and costs by the same factor will tend to emphasise the costs more. Note that his was a tentative, and weak, expectation, as we had no a priori way of knowing how the cost of walking a set distance compares to a set sucrose reward.

### Data Analysis

The entire statistical analysis code, including data handling, figure code, and analysis results, is presented in supplement ESM2. Raw data is available in supplement ESM1. All the statistical analyses were performed in R 4.1.2 (R Core Team, 2020). The packages readODS (Schutten et al., 2020) and reshape2 (Wickham, 2007) were used to load and prepare the data. We focused on two measures: the binomial choice at the last experimental visit, and the number of pheromone depositions during the training visits for the different options.

To analyse the former, we employed generalized mixed effect models with a binomial distribution using the package glmmTMB (Brooks et al., 2017; Magnusson et al., 2020). In every experiment, we included as predictors the choice order (first, second or third visit to the Y-maze) and decision line (initial decision, passed the first 2cm line; or final decision, reached the end of the arm). The goodness of fit was evaluated with the package “DHARMa” (Hartig, 2018). We performed an analysis of deviance to observe the effect of the predictors using the package car (Fox and Weisberg, 2011), and then performed Bonferroni-corrected post-hoc analysis on predictors that have an effect using the package emmeans (Lenth et al., 2020).

For pheromone deposition we followed the same procedure. We employed GLMM with a Poisson error structure, varied into a Tweedie error structure when DHARMa testing suggested that as appropriate. In the pilot, we included direction (to the drop or back to the nest) and visit length (short or long) as predictors. For the other three conditions, we included runway section (nearest to the nest, middle, nearest to the end) and treatment of the visit (‘Bundled’, ‘Segregated All’, ‘Segregated Reward’). We then followed up with analyses of deviance and post-hoc analyses as well.

In the pilot experiment, we expected the ants to deposit more pheromone on the long runway, independently of their preference. All things being equal, a triple-length runway will offer triple the pheromone deposition time. To control for this bias, we multiplied the observed pheromone deposited on the short runway by three. This may not be wholly appropriate, however, if for example pheromone is not deposited evenly throughout the path length. Indeed, *L. niger* are reported to deposit more pheromone nearer to the food source (Beckers et al., 1992). We thus advise caution in the interpretation of the pheromone data from the pilot experiment. This is not a problem for the three experimental conditions as the runway length remains fixed.

After analysis, the data was then passed onto a Python 3 (Van Rossum and Drake, 2009) environment using the package reticulate (Ushey et al., 2021), to produce graphs. To achieve this, we used the libraries pandas (Jeff Reback et al., 2020), numpy (Oliphant, 2006; van der Walt et al., 2011), matplotlib (Hunter, 2007) and seaborn (Waskom et al., 2017).

Examining the results of the aforementioned model, we discovered a possible contrast effect, as depending on the condition, the amount of pheromone deposited for the same treatment changed abruptly. It is indeed crucial to consider how the first experience of ants can have ripple effect on its subsequent decisions (De Agrò et al., 2021). To further examine this effect we remodeled the data including the first experienced treatment as a factor. We will present the results of the model that included and that did not include the first experience separately. The full analysis, and all the raw data, can be found in the supplements.

## Results

### Pilot Experiment – Cost of travelled distance

In the pilot experiment, the ants were asked to choose between two odours: one associated with a short runway, and the other associated with a long one. We observed a 90% probability of the ants choosing the short-associated odour when encountering the Y-maze for the first time, significantly higher than chance level (GLMM post-hoc: prob.=0.896, SE=0.056, t=3.589, p=0.0014). The probability quickly dropped to chance level for the two subsequent visits (visit 2: prob.=0.547, SE=0.124, t=0.376, p=1; visit 3: prob.=0.484, SE=0.123, t=-0.129, p=1). This is generally to be expected in this type of experiment, as the lack of a reward and manipulation easily disrupts the ant decision.

The pheromone deposition analysis confirms this pattern, as indeed ants deposit double the amount of pheromone per unit length on the short runway over the long one (GLMM post-hoc ratio=2, SE=0.231, t=5.973, p<0.0001).

### Main experiment – Bundling vs Segregation

#### Condition 1) ‘Segregated Reward’ Vs ‘Bundled’

In this experiment, we expected ants to prefer the ‘Segregated Reward’ treatment over the ‘Bundled’ one due to bundling.

We observed an odd difference between the three subsequent tests on the Y-maze (GLMM ANODA, chi-square=9.5744, DF=2, p=0.0083). Specifically, the ants showed no significant preference in the first visit (GLMM post-hoc: prob.=0.516, SE=0.0693, t=0.227, p=1) nor in the third (prob.=0.486, SE=0.0693, t=-0.195, p=1). However, they significantly preferred the segregated option in the second visit (prob.=0.729, SE=0.0.0596, t=3.279, p=0.0036). We consider this a false positive (see discussion).

For the pheromone deposition (Figure 3), we observed a difference between the treatments (GLMM ANODA, chi-square=6.82, DF=1, p=0.009). Here, we also observed a difference between the three sections (chi-square=57.38, DF=2, p<0.0001), but there was no effect of the interaction between the two other predictors (chi-square=0.6293, DF=2, p=0.73). Specifically, there was a non-significant trend of more pheromone deposited for the ‘Segregated Reward’ option than for the ‘Bundled’ one (GLMM post-hoc: ratio=0.835, SE=0.0618, t=-2.438, p=0.0597). The ants deposited overall more pheromone on the section of the runway nearest the drop in respect to the second (ratio=1.548, SE=0.1324, t=5.108, p<0.0001) or third (ratio=1.881, SE=0.1676, t=7.091, p<0.0001) section.

**Figure 3:**
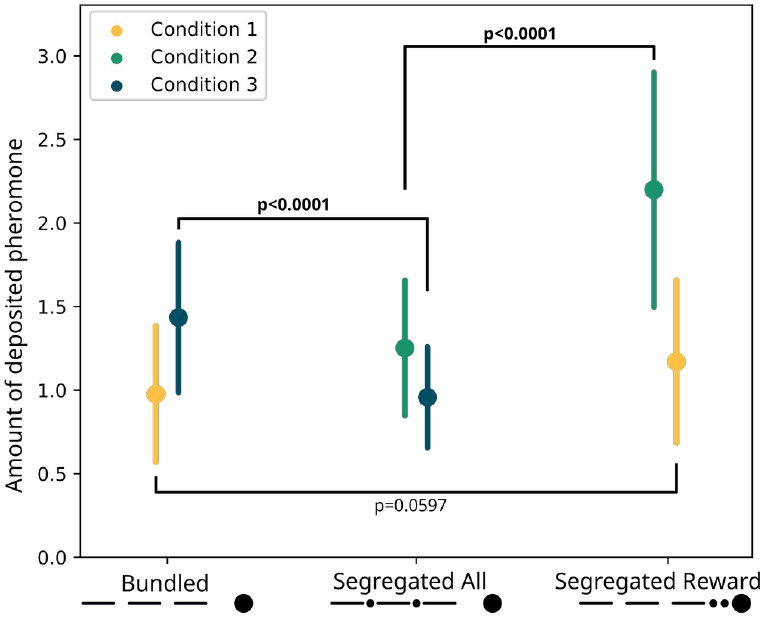
Modelled pheromone deposition for each treatment, across the three condition. Y-axis: amount of deposited pheromone per runway section. Error bars represent standard error. In yellow, pheromone deposited in the ‘Bundled” vs ‘Segregated Reward” condition (n=40). The two treatments are not significantly different from each other (p=0.0597). In green, pheromone deposited in the ‘Segregated All” vs ‘Segregated Reward” condition (n=40). The two treatments are significantly different from each other (p<0.0001). In blue, pheromone deposited in the ‘Segregated All” vs ‘Bundeled” condition (n=40). The two treatments are significantly different from each other (p<0.0001).

Regarding the effect of the first encountered treatment (Figure 4), we observed that when the ‘Segregated Reward’ treatment was encountered first, the ant significantly preferred it to the ‘Bundled’ option (GLMM post-hoc: ratio=0.652, SE=0.0618, t=-4.601, p<0.0001) section. Instead, when they encountered the ‘Bundled’ treatment first the ants showed no preference (GLMM post-hoc: ratio=1.127, SE=0.1297, t=1.041, p=1).

**Figure 4:**
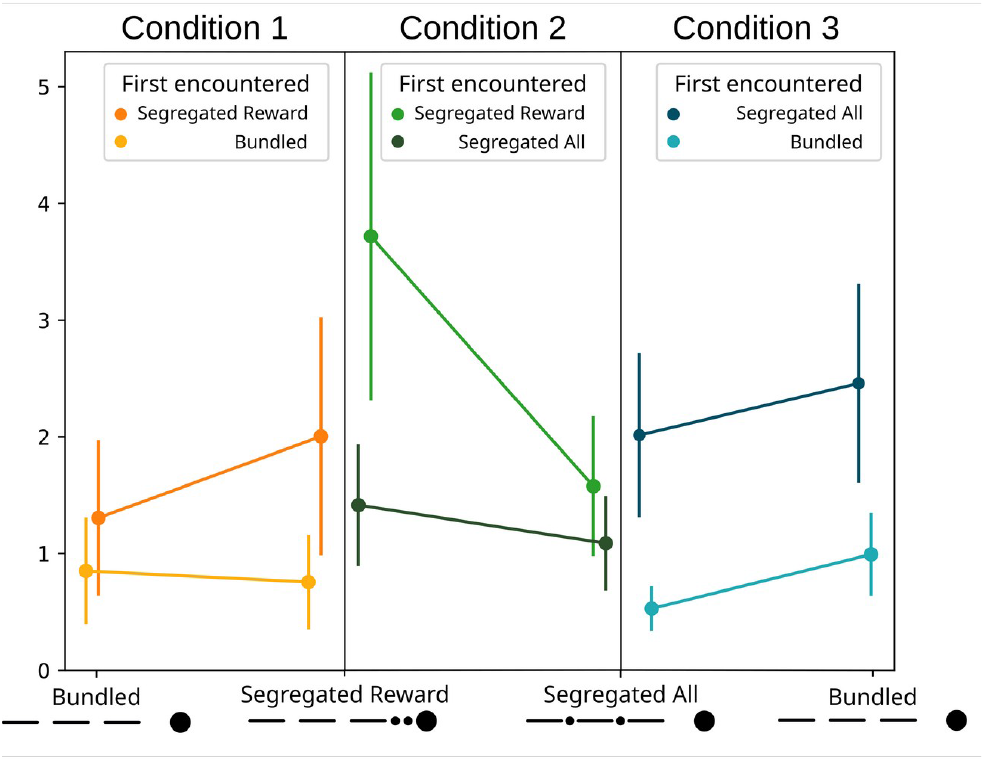
Modelled pheromone deposition for each treatment, across the three condition, including the influence of the first experienced option. Y-axis: amount of deposited pheromone per runway section. Error bars represent standard error. When ‘Segregated Reward’ is encountered first, both in condition 1 and 2, the overall pheromone deposited is higher in respect to the ‘Bundled’ (p<0.0001) and to the segregated all option (p<0.0001). In condition 3, when the ‘Bundled’ option is encountered first it is preferred over the ‘Segregated All’ option (p<0.0001). When ‘Segregated Reward’ is instead not encountered first both in condition 1 and 2, ants deposited the same amount of pheromone for both options (condition 1: p=1, condition 2: p=0.1572). Similarly, in condition 3, ants deposited the same amount of pheromone for both options when ‘Segregated All’ treatment was experienced first (p=0.2555).

#### Condition 2) ‘Segregated Reward’ vs ‘Segregated All’

In this experiment we expected the ‘Segregated Reward’ treatment to be preferred over the ‘Segregated All’ treatment.

We found no difference between subsequent Y-maze visits (GLMM ANODA: chi-square=3.1754, DF=2, p=0.2044), and we observed no overall significant preference in the Y-maze test (GLMM post-hoc: prob.=0.524, SE=0.06, t=0.402, p=0.688).

Regarding pheromone deposition (Figure 3), we observed a difference between the treatments (GLMM ANODA, chi-square=57.8966, DF=1, p<0.0001) and a difference between the three sections (chi-square=9.7448, DF=2, p=0.0076). We also observed an almost significant effect of the interaction (chi-square=5.9719, DF=2, p=0.0504). Specifically, the ants deposited more pheromone for the ‘Segregated Reward’ option compared to the segregated cost one (GLMM post-hoc: ratio=0.569, SE=0.0426, t=-7.529, p<0.0001). In the ‘Segregated Reward’ visits, but not the ‘Segregated All’ visits, the ants deposited more pheromone on the section of the runway nearest the drop relative to the furthest one (ratio=1.56, SE=0.1758, t=3.95, p=0.0008).

Regarding the effect of the first encountered treatment (Figure 4), we observed that when the ‘Segregated Reward’ treatment was encountered first, the ant significantly preferred it to the ‘Segregated All’ option (GLMM post-hoc: ratio=0.434, SE=0.0412, t=-8.831, p<0.0001) section. However, when they encountered the ‘Segregated All’ treatment first the ants showed no preference (GLMM post-hoc: ratio=0.769, SE=0.0907 t=-2.227, p=0.1572).

#### Condition 3) ‘Segregated All’ Vs ‘Bundled’

In this experiment, ‘Bundled’ treatment was expected to be preferred over the ‘Segregated All’ one.

The ants showed no overall significant preference for either treatment (GLMM post-hoc prob.=0.466, SE=0.122, t=-0.279, p=0.7803).

However, we saw large differences in pheromone deposition between the treatments (Figure 3). We found an effect of the two treatments (GLMM ANODA, chi-square=26.52, DF=1, p<0.0001), of the three runway sections (chi-square=13.43, DF=2, p=0.0012) and of the interaction between those (chi-square=8.16, DF=2, p=0.0169). Specifically, ants deposited more pheromone in the ‘Bundled’ visits than in the segregated cost ones (GLMM post-hoc: ratio=1.5, SE=0.123, t=4.91, p<0.0001). Moreover, in the ‘Bundled’ visits, they deposited more pheromone on the runway section nearest the drop relative to the respect to the middle section (ratio=1.538, SE=0.195, t=3.4, p=0.007) or the section nearest the bridge (ratio=1.729, SE=0.223, t=4.243, p=0.0002). This pattern was not present in the segregated cost visits. (first vs second section: ratio=1.065, SE=0.163, t=0.412, p=1; first vs third section: ratio=0.995, SE=0.150, t=-0.032, p=1).

Regarding the effect of the first encountered treatment (Figure 4), we observed that when the ‘Bundled’ treatment was encountered first, the ant significantly preferred it to the ‘Segregated All’ option (GLMM post-hoc: ratio=0.533, SE=0.0734, t=-4.571, p<0.0001) section. Instead, when they encountered the ‘Segregated All’ treatment first the ants showed no preference (GLMM post-hoc: ratio=0.820, SE=0.0803 t=-2.031, p=0.2555).

## Discussion

Bundling and segregation treatment strongly affected the ants’ perceived value, but not their choices. Analysing the binary choices of the ants in the main study, no consistent preference was found. While ants significantly preferred the segregated reward over the bundled reward in their second of three trials, we can conceive of no plausible biological or psychological reason for this to be so, and interpret this as a false positive. Thus, there is no evidence that bundling or segregation affect how much these insects “want” the reward.

However, the pattern of pheromone deposition, which correlates strongly with perceived food quality in these and other ants (Beckers et al., 1993; Czaczkes et al., 2018b; Jackson and Châline, 2007; Wendt et al., 2019), was fully in line with our predictions: ants deposited most pheromone when the costs were bundled and rewards segregated, and a lot less in the other two treatments (Figure 3). Our weaker prediction of ‘Bundled’ being “liked” more than ‘Segregated All’ was also supported. Thus, while bundling and segregation does not affect “wanting” in these insects, it seems to affect “liking”, directly in line with predictions from classical behavioural economic and perception research.

However, the pattern is somewhat more complicated, as we saw large differences in pheromone deposition for the same treatment, depending on the treatments they were paired with. For example, while ‘Segregated Rewards’ are recruited to almost twice as strongly as ‘Segregated All’ when they are paired, ‘Segregated Rewards’ are recruited to much more weakly when paired with the ‘Bundled’ treatment (see Figure 3). A similar pattern is seen with ‘Bundled’, where recruitment is stronger when paired with ‘Segregated All’ and weaker when paired with ‘Segregated Reward’. This pattern is further complicated by a strong effect of first encountered treatment (see Figure 4). One possibility is that these differences arise due to a series of contrast effects.

Notably, the preference for ‘Bundled’ over ‘Segregated All’, and the preference for ‘Segregated Reward’ over ‘Segregated All’, are very clear. By contrast, the preference for ‘Segregated Reward’ over ‘Bundled’ is less clear, appearing only when disentangled from the ‘first encountered treatment’ effect. This seems counter-intuitive, as the ‘Bundled’ and ‘Segregated Reward’ options only differ in the boosting of a gain. Moreover, the contrast of ‘Segregated All’ versus ‘Bundled’, for which we had no strong a-priori prediction as we had no idea how gains and losses are weighted, shows a very clear difference. These results are consistent with a very strong segregation effect for losses, and a very mild one, if any, for gains. Assuming this is the case, the ‘Bundled’ and the ‘Segregated Reward’ treatments will be considered almost identical, bar the mild preference for ‘Segregated Reward’. In the contrast between ‘Segregated All’ and ‘Bundled’, the preference for bundled becomes clear, as the segregation of losses drives the preference completely. The preference for ‘Segregated Reward’ over ‘Segregated All’ remains equally clear, as the two options only differ in the realm of losses. Overlaid onto this pattern is the ‘first encountered treatment’ effect: in every condition the effects strengthen when the preferred option is presented first, and drops to chance level when presented second. This suggest that the bias generated through the bundling process has a similar strength to the preference for the first experienced odour (Oberhauser et al., in revision), and thus they appear to counterbalance each other.

Bundling and segregation of options to modify perceived value in insects may have ecological implications, especially in plant-pollinator interactions. If, by segregation, the perceived value of a reward can be increased, plants may be selected to split rewards amongst multiple smaller flowers in an inflorescence, or flowers on a plant. Similarly, they may be selected to attempt to bundle costs, favouring multi-flower inflorescences, allowing insects to walk between flowers, over multiple small, separate flowers.

The finding that bundling and segregation affect ant value perception adds this behavioural economic effect to several others which have also been shown to affect perceived value in insects, including decoy effects (Sasaki and Pratt, 2011; Shafir et al., 2002; Tan et al., 2015), invested effort increasing perceived value (Czaczkes et al., 2018a), relative value perception (Bitterman, 1976; Couvillon and Bitterman, 1984; Wendt et al., 2019), and labelling effects (Hemingway and Muth, 2022; Wendt and Czaczkes, 2020). It is becoming clear that the underlying patterns driving value perception in insects are in many ways parallel those of humans. This implies either an extremely early evolutionary origin of shared perceptual mechanisms resulting in shared psychophysical laws, or convergent evolution on similarly-effective systems. The finding of a bundling and segregation effect, alongside the way in which insects respond to differences between experienced and expected rewards, strongly implies that insects share the same broad value function shape as humans (see Figure 1). Thus, we would predict that any other behavioural economic effects in humans which arise from this value function, to also be present in insects. Still missing is a demonstration of an inflection point –that, like in humans, losses loom larger than gains for insects. That our results imply bundling to more strongly influence losses than gain (see above) suggests this is the case, but formal testing will be required.

Our experiment revealed a mismatch between instantaneous evaluation of an option (pheromone deposition) and choice. We hypothesize that the former reflects “liking”, while the latter reflects “wanting”, and that these are separate functions in insects. “Liking” and “wanting” in vertebrates are neurologically separate, and respond differently in a variety of situations (Leyton et al., 2002; Treit and Berridge, 1990). Moreover, in humans, instantaneous hedonic ratings (how much subjects like something right after experiencing it) can be highly divergent from ratings given after cognitive processing (e.g. Wilson and Schooler, 1991), and this may well be true in insects as well. To our knowledge, the current study is the first to distinguish “liking” and “wanting” in an insect. Pheromone deposition behaviour in ants offers a unique window into the mind of an insect. When travelling away from a food source, the animal must by definition be divorced from “wanting”, as it is heading away from the reward. As pheromone deposition correlates very strongly with perceived reward value (Beckers et al., 1993; Breed et al., 1987; Crawford and Rissing, 1983; Jackson and Châline, 2007; Wendt et al., 2019), we consider it to be a good metric of “liking” which is, at least on the return to the nest, disassociated from “wanting”. While pheromone deposition (“liking”) very often co-vary with choice behaviour (“wanting”) (Czaczkes et al., 2018a; Wendt et al., 2020, 2019; Wendt and Czaczkes, 2020), the results of the current experiment empirically demonstrates that they can be dissociated. However, being such an important behaviour, pheromone deposition is also carefully modulated in other situations as well. For example, ants increase pheromone deposition after correcting a navigational error (Czaczkes and Heinze, 2015) or when foraging at low light levels (Jones et al., 2019), and decrease pheromone deposition on busy trails (Czaczkes et al., 2014, 2013), when returning from occupied food sources (Wendt et al., 2020), or when the food is associated with danger (Nonacs, 1990). This makes using pheromone deposition as a pure measuring of “liking” in all situations challenging. Most importantly, *L. niger* almost never deposit pheromone if they have not drunk to satiation (Mailleux et al., 2000), and so we can only observe this measure of “liking” for a non-random subset of ants which “want” the food sufficiently to drink to satiation.

The “wanting” vs “liking” framework suggests that mental processes that we assume to be connected can instead be separate. Scalar Utility Theory (SUT; Kacelnik and Brito e Abreu, 1998; Rosenström et al., 2016) is an influential framework that has been developed to describe decision making under uncertainty. This theory postulates that encoding neurons, responsible for the internal representation of values (i.e. quantity, quality, delay, etc.), have logarithmically spaced sensitivities and specificities. Prospect theory also assumes a process based on a logarithmic function. Thus, both make very similar behavioural predictions, even though one concentrates on value perception, and the other on memory acquisition. The “liking” vs “wanting” distinction however, especially given the results observed in this experiment, suggests that perception and memory can be segregated process. Most experiments recording both pheromone deposition and subsequent choice use food quality (from high concentration sucrose to distasteful solutions) as the measured unit (De Agrò et al., 2021; Oberhauser and Czaczkes, 2018; Wendt et al., 2019). Logarithmic perception of quality is probably the reason mismatch a mismatch between choice and instantaneous value perception has eluded discovery. We believe that the mismatch produced in this experiment has to do with the type of cost chosen as treatment: distance traveled. As previously mentioned, our treatment seemed unable to produce a segregation effect for the reward, while showing a strong segregation effect for losses, i.e. the traveled distance. It is reasonable to assume that ants perceive value logarithmically, in accordance to Prospect Theory (De Agrò et al., 2021). In other words, overall enjoyableness of the experience (i.e. “liking”) is affected by the artifacts of a log curve, segregation effect included (see Figure 1). Being produced instantaneously, pheromone deposition is likely linked to this information stream. However, when registering distances in memory ants need to be extremely precise. They possess an internal step counter and a visual odometer, that allows them to judge distances and return successfully to the nest (Narendra, 2007; Wittlinger et al., 2006). Encoding steps logarithmically would be absolutely insufficient for accurate homing, given the need to already cope with errors introduced in the path integration process (Merkle et al., 2006; Merkle and Wehner, 2010; Müller and Wehner, 1988; Schwarz et al., 2011). For this, a linear, 1:1 correspondence is required. It is possible that distance is memorized linearly and in a separated stream from the hedonistic perception.

When asked to choose, the ant can only compare the two memories of distances, which due to their linear nature are immune to the segregation effect (Figure 5). If the ant odometer truly is a rare example of linear perception, it would be an invaluable system for investigating information processing in insects, as is allows the roles of perception and post-perceptual processing to be disentangled.

**Figure 5:**
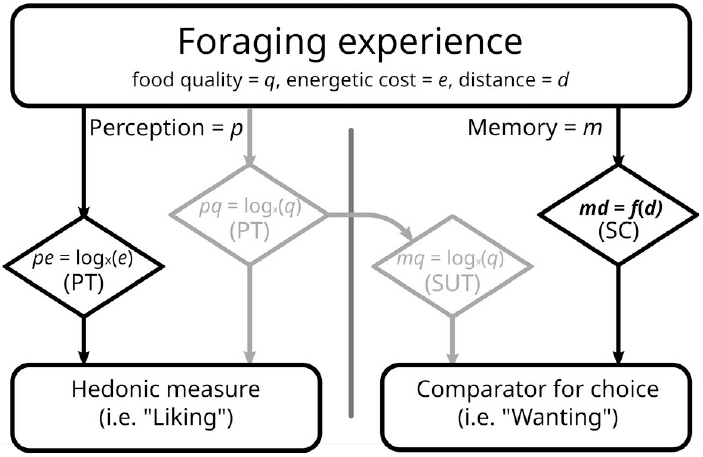
Proposed model of liking vs wanting in light of economic theories. Percetion can be connected to the formed memory, but not necessarily. During a foraging bout, the ant perceive (p) gains (food quality, q) and losses (energy spent to reach, e), according to Prospect Theory (PT). Food quality seem to be registered into memory (m) in to the same scale, congruently with Scalar Utility Theory (SUT). Distance travelled (d) represent a special case, as it requires precise memory in the context of ant navigation. As such, it may possess a dedicated, direct, and linear memorization circuit (md), like the Step Counter (SC). In our experiment, we failed to imprint a segregation effect into rewards (grayed out boxes. See results and discussion), and as such all our options were equal in this realm. With costs perceived logarithmically, but memorized linearly, we would expect the results observed in this experiment.

Our work also demonstrates empirically, for the first time, that individual ants prefer near to distant food sources. Ants also recruited significantly more to closer food source, as reported in this and other ant species as well (Devigne and Detrain, 2006; Fewell et al., 1992), implying both increased “liking” and increased “wanting”. However, this result is misleading, as *Lasius niger* deposit much more pheromone in the immediate vicinity of the reward (Devigne and Detrain, 2006). Thus, while ants clearly “want” nearer food sources, we cannot say whether they “like” them more.

This study offers several ‘firsts’ – first, almost trivially, it is the first demonstration of ant individual preference for nearer food. More importantly, it is the first demonstration of the bundling and segregation effect influencing value perception in an insect. This further supports the case that insect and human value functions are surprisingly similar. Finally, and perhaps most importantly, this is the first introduction of the concepts of “liking” and “wanting” in an invertebrate, and the first empirical support for their being separate, using ant trail pheromone deposition as a measure of “liking”, and choice as a measure of “wanting”. Ant trail pheromone deposition behaviour is a useful window into the mind of an insect. However, we hope that this demonstration will encourage other researchers to take other approaches to disentangle “wanting” from “liking” in insects. It took over a decade for the liking/wanting distinction to move from rats (Berridge et al., 1989) to humans e.g. (Reviewed in Berridge, 2018; Brauer and De Wit, 1997; Leyton et al., 2002). Twenty years later, the move to invertebrates is long overdue.

## Acknowledgements

TJC was supported by a Heisenberg fellowship from the Deutsche Forschungsgemeinschaft (CZ 237 / 4-1). MDA was funded by University of Regensburg ‘Anreizsystem’ funding to TJC.

## Competing Interests

The authors declare no competing interests.

## Notes

### Competing Interest Statement

The authors have declared no competing interest.

